# Improved prediction of stabilizing mutations in proteins by incorporation of mutational effects on ligand binding

**DOI:** 10.1101/2024.04.11.589149

**Authors:** Srivarshini Ganesan, Nidhi Mittal, Akash Bhat, Rachna S. Adiga, Ananthakrishnan Ganesan, Deepesh Nagarajan, Raghavan Varadarajan

## Abstract

While many computational methods accurately predict destabilizing mutations, identifying stabilizing mutations has remained a challenge, due to their relative rarity. We tested ΔΔG^0^ predictions from computational predictors such as Rosetta, ThermoMPNN, RaSP, and DeepDDG, using eighty-two mutants of the bacterial toxin CcdB as a test case. On this dataset, the best computational predictor is ThermoMPNN which identifies stabilizing mutations with a precision of 68%. However, the average increase in T_m_ for these predicted mutations was only 1°C for CcdB, and predictions were poorer for a more challenging target, influenza neuraminidase. Using data from multiple previously described yeast surface display libraries and *in vitro* thermal stability measurements, we trained logistic regression models to identify stabilizing mutations with a precision of 90% and an average increase in T_m_ of 3°C for CcdB. When such libraries contain a population of mutants with significantly enhanced binding relative to the corresponding wild type, there is no benefit in using computational predictors. It is then possible to predict stabilizing mutations without any training, simply by examining the distribution of mutational binding scores. This avoids laborious steps of *in vitro* expression, purification, and stability characterization. When this is not the case, combining data from computational predictors with high-throughput experimental binding data enhances the prediction of stabilizing mutations. However, this requires training on stability data measured *in vitro* with known stabilized mutants. It is thus feasible to predict stabilizing mutations rapidly and accurately for any system of interest that can be subjected to a binding selection or screen.

## 1 Introduction

Protein-based therapeutics, vaccines, and enzymes have many important biomedical and biotechnological^1,2^ applications. In most cases, it is desirable to stabilize the protein to prevent its unfolding or aggregation in non-native environments and improve product efficacy^3–6^. Experimental identification of stabilizing mutations is often time-consuming and expensive. Exhaustive exploration of mutational space through experiments can be technically challenging, especially for large proteins. This is in part because most mutational screens use ligand binding or some other measure of protein activity as their readout. This results in the selection of better binders/more active mutants which are not necessarily more stable.

In an effort to overcome this lacuna, we have previously developed saturation suppressor methodology^7,8^ in which the mutational library is constructed in the background of a destabilizing mutation (termed as a parent inactivating mutation or PIM), typically positioned at a buried site. When introduced into the wild-type background, this mutation causes misfolding, and/or aggregation or degradation, leading to consequent loss of activity. Therefore, this mutant creates a possibility to select second-site suppressors that lead to the regaining of function by introducing the PIM into every member of a saturation library of the protein of interest and screening for clones that are properly folded as assessed by their activity or ability to bind a conformation-specific ligand. When such suppressors are located far from the original destabilizing mutation (distal suppressors), they are typically able to suppress deleterious effects of a variety of destabilizing mutations located at different structural locations and are therefore classified as global suppressors. Such suppressors typically enhance protein stability *in vitro* and foldability and yield *in vivo*, even in the wild-type context^7–9^. In contrast, suppressors in physical contact with the destabilizing mutation (local or proximal suppressors) are generally allele-specific and are often destabilizing in the wildtype context, offering a facile way to distinguish between global and proximal suppressors in the absence of any structural information^8^. We have used this approach to isolate a large number of stabilizing mutations of the bacterial protein CcdB^7^ as well as of the receptor binding domain of the Spike protein of SARS-CoV-2^10^.

CcdB is a homodimeric bacterial toxin that binds to DNA gyrase and causes bacterial cell death. CcdB is an ideal system for understanding the effects of mutation on stability due to its small size (101 amino acids), facile phenotypic readout, and extensive prior experimental characterization^11–13^. We screened a number of saturation suppressor libraries of yeast surface displayed CcdB for binding to the DNA Gyrase fragment, GyrA. In this approach, binding is quantitated by estimating the mean fluorescence intensity of binding (MFI) of each member of the library to epitope tagged GyrA using FACS coupled to deep sequencing. CcdB mutants with significantly higher binding MFI values than the corresponding PIM are putative stabilized mutants^7^. Subsequently, ∼85 such individual mutants were expressed in E. coli, purified and the *in vitro* thermal stability was measured to validate the methodology^7^.

Many computational methods have been developed to predict the change in the Gibbs free energy of folding (ΔΔG^0^) upon point mutations^14^. Existing computational methods can be broadly classified into biophysics-based and data-driven approaches. With the success of deep learning in predicting protein 3D structures, many attempts have been made to develop models for ΔΔG^0^ prediction using related approaches, and by combining protein sequence, structure, and experimental stability data. While some state-of-the-art deep learning methods have achieved accuracies as high as 80% for predicting the value of ΔΔG^0^ for point mutations, only about 20% of the predicted stabilizing mutations were found to be stabilizing in experiments^15^. This is because the fraction of stabilizing mutations is low, both in sequence space and in the existing experimental data upon which these deep-learning models are trained^16^.

We utilized the computational stability predictors Rosetta, ThermoMPNN, RaSP, and DeepDDG^17–19^ to predict the ΔΔG^0^ for point mutations in CcdB. Rosetta is primarily a protein design suite that has been used for structures ranging from miniproteins^20^ to megadalton-scale protein complexes^21^. ThermoMPNN uses a pre-trained message-passing graph neural network that predicts the most favorable amino acid for a given 3D environment. The output is then passed through a stability prediction module that is finetuned on a large experimental stability dataset^22,23^ to predict the ΔΔG^0^ for that point mutation. RaSP uses a Convolutional Neural Network that is finetuned on Rosetta calculated ΔΔG^0^ values to predict the effect of point mutations. DeepDDG is a neural network trained on experimental data obtained from the ProTherm database^24^. It includes various geometric, sequence, and general features like the solvent-accessible surface area, backbone dihedrals, secondary structure, position-specific scoring matrix values, and amino acid type.

To further enhance the prediction of stabilizing mutations, we built logistic regression models using both the experimental mutagenesis data and computational stability predictors. When the experimental libraries contain a population of mutants with significantly enhanced binding relative to the corresponding wild type, we observed that there is no benefit in using computational predictors to predict stabilizing mutations. However, when this is not the case, combining data from computational predictors with high-throughput experimental binding data enhances the prediction of stabilizing mutations.

Additionally, we analyzed computational predictions of stabilizing mutations in a complex target, Neuraminidase (NA), a homotetrameric surface glycoprotein of the Influenza virus. NA and hemagglutinin (HA) serum inhibition titers have been identified as correlates of protection^25–27^. In contrast to natural infection, seasonal vaccination fails to induce a robust anti-NA immune response because of the low amount and poor stability of NA in current vaccine formulations^28,29^. Well-expressed and correctly folded NA antigens will likely induce protective antibody responses. Since NA and HA natural sequence variations are uncorrelated^30^, the addition of NA to existing influenza vaccines can potentially enhance vaccine efficacy. We therefore used ThermoMPNN to predict stabilizing mutations for NA and tested the predictions experimentally through cell surface expression and binding studies.

## 2 Materials and methods

### 2.1 MFI calculation for CcdB libraries

We obtained binding MFI values in the background of the wildtype CcdB^31^ and for different PIM libraries with destabilizing mutants V18G, V20G, or L36A^7^. The MFI for each mutant was normalized relative to the MFI of the respective PIM (Equation 1) or wildtype (Equation 2). The normalized MFI values were rescaled to lie between 0 and 1 (Equation 3).

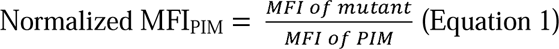

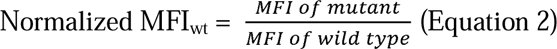

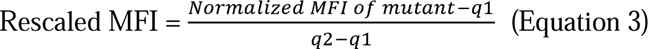

where q1 and q2 are the 5^th^ and 95^th^ percentile of the sorted normalized MFI values. The rescaled MFI value was assigned to be 1 for mutants with normalized MFI values higher than q2 and 0 for mutants with normalized MFI values lower than q1. We next calculated the average of the rescaled MFI values from the three PIM libraries (Equation 4).

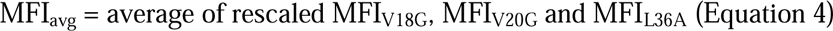

### 2.2 Obtaining Computational Predictions

The Rosetta ΔΔG^0^ predictions were obtained by performing the Rosetta software suite^32,33^ (version 3.14). Ccdb (PDB ID: 3vub) was initially subjected to all-atom energy-minimization (relax.static.linuxgccrelease) using the default energy function. Initially, 100 iterations of energy minimization were performed and the solution possessing the lowest total energy score was used as the template structure for further steps. This ensures that the structure adopts the lowest possible energetic state (ΔG_1_), as measured by the Rosetta energy function. Secondly, a given amino acid (m, from 1-101 total amino acids) was mutated to a selected identity (n, from 1-19 canonical amino acids) using fixed backbone design (fixbb.static.linuxgccrelease) using the default energy function. Fixed backbone design would also select the lowest-scoring rotamer for a given amino acid mutation. The designed mutant would then be subjected to a further 100 iterations of all-atom energy-minimization (relax.static.linuxgccrelease). The energy scores from the ten lowest scoring structures were averaged (ΔG_2_). For a given residue position, and a given mutant, the ΔΔG^0^ was simply: ΔΔG^0^ = ΔG_2_-ΔG_1_. ΔΔG^0^ was evaluated m*n times, for each residue position and each amino acid mutant.

The ThermoMPNN and RaSP predictions were obtained by running their prediction algorithms available through Google Colab (https://colab.research.google.com/drive/1OcT4eYwzxUFNlHNPk9_5uvxGNMVg3CFA, https://colab.research.google.com/github/KULL-Centre/papers/blob/main/2022/ML-ddG-Blaabjerg-et-al/RaSPLab.ipynb). For DeepDDG, the ΔΔG^0^ predictions were made using their webserver (https://protein.org.cn/ddg.html). For all the predictors, the coordinates of the CcdB homodimer were obtained from the PDB ID 3VUB^34^.

### 2.3 Metrics to evaluate the models

The precision, recall, and Matthews correlation coefficients were calculated as: Precision = TP/(TP+FP) (Equation 5)

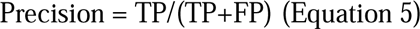

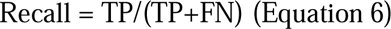

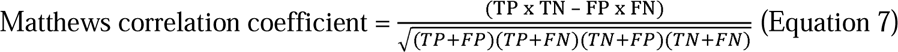

where TP, FP, and FN denote the number of True Positives, False Positives, and False Negatives, respectively.

TP: number of mutants for which predicted ΔT_m_ > 0 and experimental ΔT_m_ > 0

TN: number of mutants for which predicted ΔT_m_ < 0 and experimental ΔT_m_ < 0

FP: number of mutants for which predicted ΔT_m_ > 0 and experimental ΔT_m_ < 0

FN: number of mutants for which predicted ΔT_m_ < 0 and experimental ΔT_m_ > 0

The average ΔT_m_ was used to evaluate the computational predictors on their prediction of stabilizing mutations and was calculated as follows. We randomly sampled 20, 40, 60, and 74 mutants from the experimentally characterized set of 82 mutants, which correspond to 25%, 50%, 75%, and 90%, respectively, of the experimental data. For each of these chosen sets of mutants, we calculated the average experimental ΔT_m_ for the subset of mutations predicted to be stabilizing by each of the computational ΔΔG^0^ predictors. We repeated the above calculation 20 times and analyzed the distribution of the 20 average ΔT_m_ values (Figure 4A).

In each iteration, the average ΔT_m_ was calculated for the predicted stabilizing mutations as:

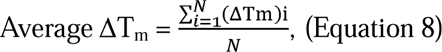

where N is the number of predicted stabilizing mutations in that iteration.

We calculated the Z-scores for all mutations in a library using the rescaled MFI values:

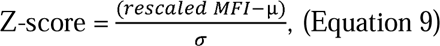

where µ and σ are the mean and the standard deviation of the rescaled MFI values.

### 2.4 Development of models for predicting stabilizing mutations

The logistic regression model is a simple classification tool that is efficient for linearly separable classes. Univariate logistic regression models have two parameters, β(0) and β(1), and can be represented as y = {3(0) + {3(1) X, where X is the feature on which the model is trained and y is the assigned class. In our training and testing sets, y = 0 indicates destabilizing mutants, and y = 1 indicates that the mutation is stabilizing. Bivariate logistic regression models have an extra parameter corresponding to the additional feature on which the model is trained. This is represented by y = {3(0) + {3(1)X(1) + {3(2)X(2) where X(1) and X(2) are the two features used to train the model. During training, parameters β(0), β(1), and β(2) are learned and then used to make predictions on the test set.

Since our models have to learn two or three parameters, the corresponding training sets must ideally have at least 20 points^35^. This corresponds to about 25% of our experimental dataset. Therefore, we trained models with 25% of the data and tested them on the remaining 75%. To understand the effect of training on a larger set, we also designed another set of models by training on 50% of the data and tested them on the remaining 50%. The process of splitting the data and training and testing the model was repeated 20 times to prevent any bias that could arise from the way the data was split. The distribution of the precision, recall, and average ΔT_m_ values of the predicted stabilizing mutations were analyzed.

### 2.5 Cloning and cell surface display of NA constructs

To further assess the performance of ThermoMPNN, we used it to predict stabilizing mutants for influenza NA from A/Wisconsin/67/2022 (H1N1). The structure of NA from this isolate was modeled using AlphaFold2^36^

(https://colab.research.google.com/github/sokrypton/ColabFold/blob/main/AlphaFold2.ipynb). The modeled structure had an RMSD of 0.77Å with its structural homolog deposited in the PDB (PDB ID: 7U2Q^37^). The viral RNA from A/Wisconsin/67/2022 (H1N1) influenza virus was extracted and cDNA encoding the full-length NA was cloned into the pcDNA3.4 vector and sequenced. To generate the ThermoMPNN predicted stabilized mutants of NA, the Neuraminidase gene was amplified via PCR using mutagenic primers followed by an overlap PCR utilizing primers incorporating XbaI (5’) and HindIII (3’) sites. The resultant product was then digested and cloned into the pcDNA3.4 vector between the above sites.

HEK293T cells were cultured in Dulbecco’s modified Eagle’s medium (DMEM) supplemented with 10% fetal bovine serum (FBS) and 1% Penstrep antibiotic. Transfections were performed using lipofectamine with 5 µg of plasmid DNA. Cells were harvested after 24 hours post-transfection and were rinsed thrice with PBS buffer containing 3% bovine serum albumin (BSA). HEK-293 cells expressing various full-length NA constructs were incubated with NA-specific antibodies (1G01, 2H08, and FNI17) at a concentration of 500 nM for 30 minutes at 4°C. After washing thrice, the cells were incubated with anti-human Alexa Fluor 633 (diluted 1:1200) for 20 minutes at 4°C. Following additional washes, NA cell surface expression in samples was analyzed using a BD Aria III instrument. Mean fluorescence intensity (MFI) was determined by subtracting the MFI of HEK-293 cells expressing neuraminidase from the MFI of HEK cells alone. The fold change in MFI was calculated as the ratio of the MFI of the mutant to the MFI of the wild type.

## 3 Results and Discussion

### 3.1 Correlation of mutant stability with mutagenesis data

We have previously experimentally characterized the thermal stabilities of a large number of CcdB mutants, both stabilizing and destabilizing^8,12,31^. We chose 82 mutants that had ΔT_m_ in the range of −7 to 10°C, with nearly equal numbers of stabilizing (38 mutants with ΔT_m_ > 0) and destabilizing (44 mutants with ΔT_m_ < 0) mutants. The mean fluorescence intensities of binding (MFI_seq_ (bind)) were obtained from two recent studies^7,31^. The MFI values of mutants introduced in the background of the wild type are henceforth denoted as MFI_wt_^31^. The MFI values of mutations introduced in the background of a parent inactivating mutation (PIM) are denoted as MFI_PIM_^7^. The destabilizing mutants V18G, V20G, and L36A were chosen as the PIMs. These were all located at buried positions in the CcdB homodimer (Figure 1).

**Figure 1.**
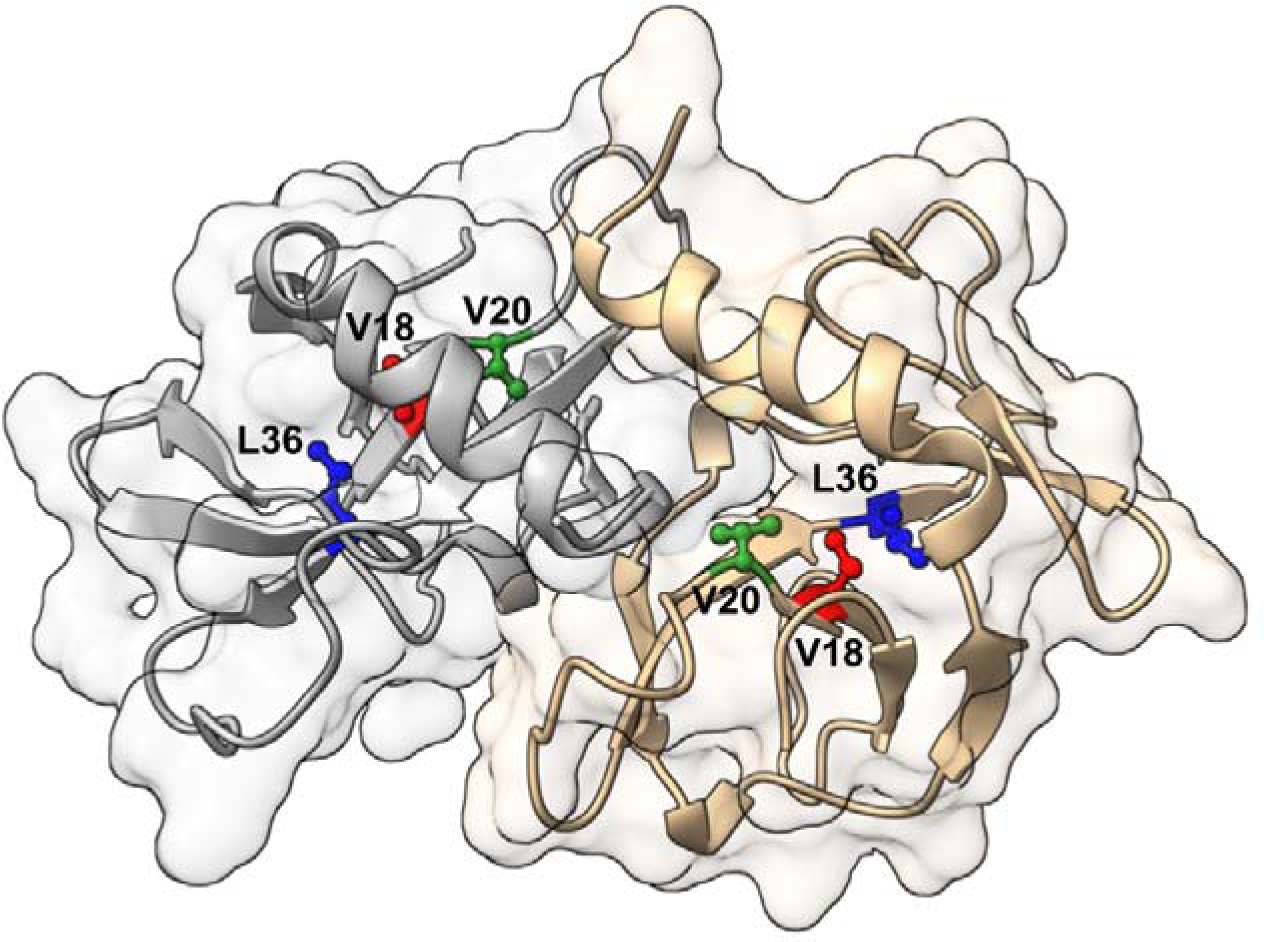
Locations of the parent inactivating mutants in the CcdB homodimer. Buried residues V18, V20, and L36 were chosen for introducing destabilizing mutations in the CcdB protein. Three saturation mutagenesis libraries were constructed in the background of each of V18G, V20G, and L36A mutations, and screened to isolate suppressors^7^.

The normalized MFI of a mutant was calculated by taking the ratio of the MFI of the mutant and the MFI of the PIM or the wild type (Equations 1 & 2). The normalized MFI values were rescaled to lie between 0 and 1 for each library (Equation 3). The rescaled values were used to calculate the average of the MFI values from the three PIM libraries (Equation 4). The rescaled MFI from each library (wildtype, V18G, V20G, and L36A), the average MFI, ΔT_m,_ and ΔΔG^0^ predicted by the computational predictors are tabulated for each of the 82 mutants and this data is available in the supplementary material. The correlation of the average MFI with experimentally measured values of the change in thermal stability (ΔT_m_) was 0.76 (Figure 2).

**Figure 2.**
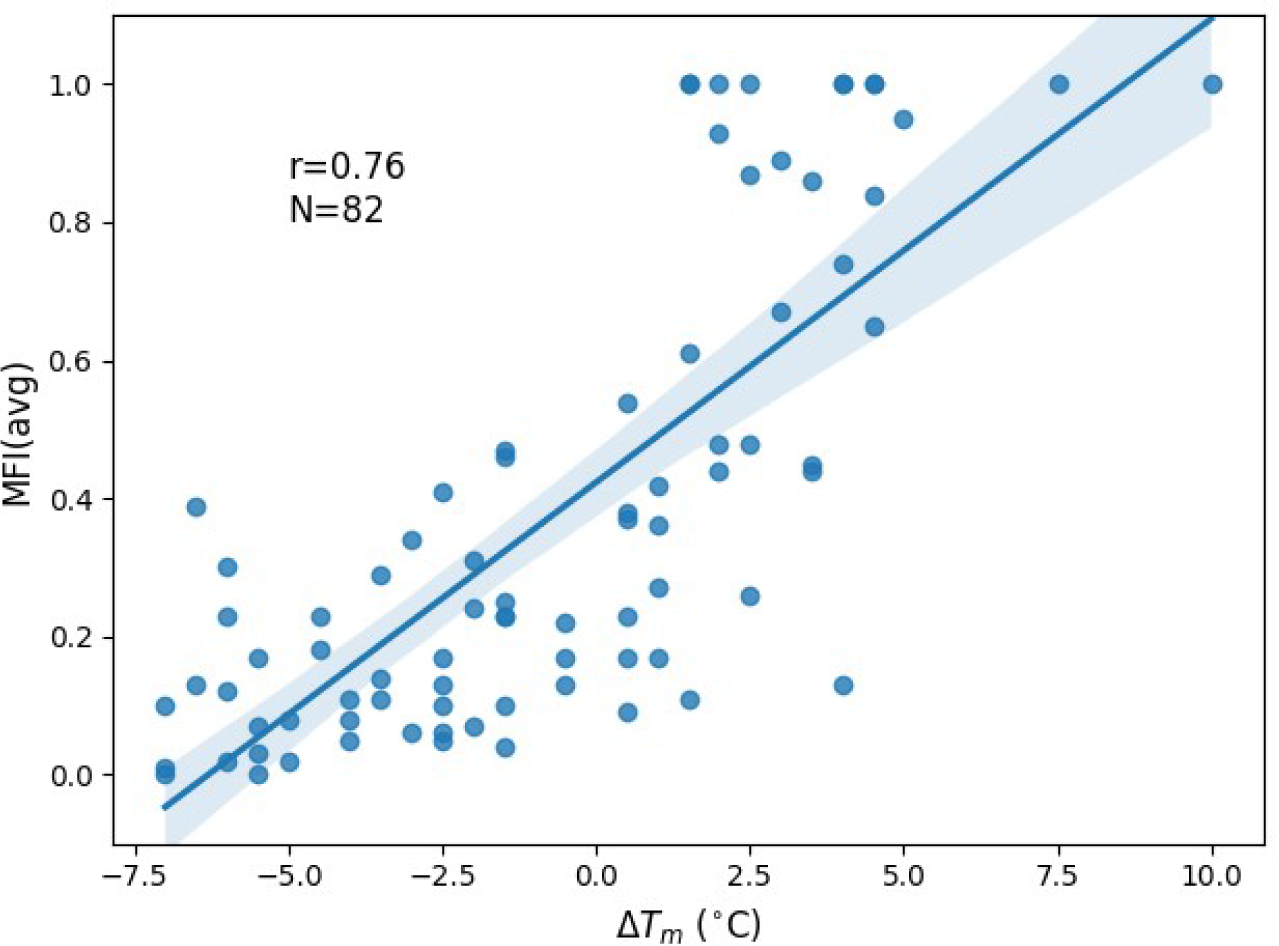
Correlation of change in mutant thermal stability (ΔT_m_) with MFI_avg_. The correlation of MFI_avg_ (Equation 4) of the 82 mutants with ΔT_m_ has a Spearman rank correlation coefficient of 0.76. We have previously shown that ΔT_m_ is linearly correlated with the change in free energy of unfolding ΔΔG^0^ ^12^.

### 3.2 Precision, recall, and correlation values for computational predictors

Several different computational stability predictors were used to predict the effect of point mutations on the stability of CcdB. ΔG^0^ represents the free energy of folding of the protein and ΔΔG^0^ = ΔG^0^_(mutant)_ – ΔG^0^_(wt)_. A mutant that is more stable than the wild type (wt) will have ΔΔG^0^ < 0. The precision, recall, and Matthews correlation coefficient (MCC) of the predicted ΔΔG^0^ values were evaluated using the experimentally determined thermal stability data (Equations 5-7). A negative value of ΔΔG^0^ will result in a positive ΔT_m_, while a positive value of ΔΔG^0^ will result in a negative ΔT_m_. ThermoMPNN had the highest precision of 0.68 followed by 0.53 for DeepDDG, 0.5 for RaSP, and 0.49 for Rosetta (Figure 3). Rosetta had the highest recall of 0.92 followed by ThermoMPNN, RaSP, and DeepDDG. The MCC was 0.27 for ThermoMPNN and 0.15 for Rosetta. For experimentalists who aim to use computational predictions to downselect mutants for detailed biophysical characterization, precision is more important than recall.

**Figure 3.**
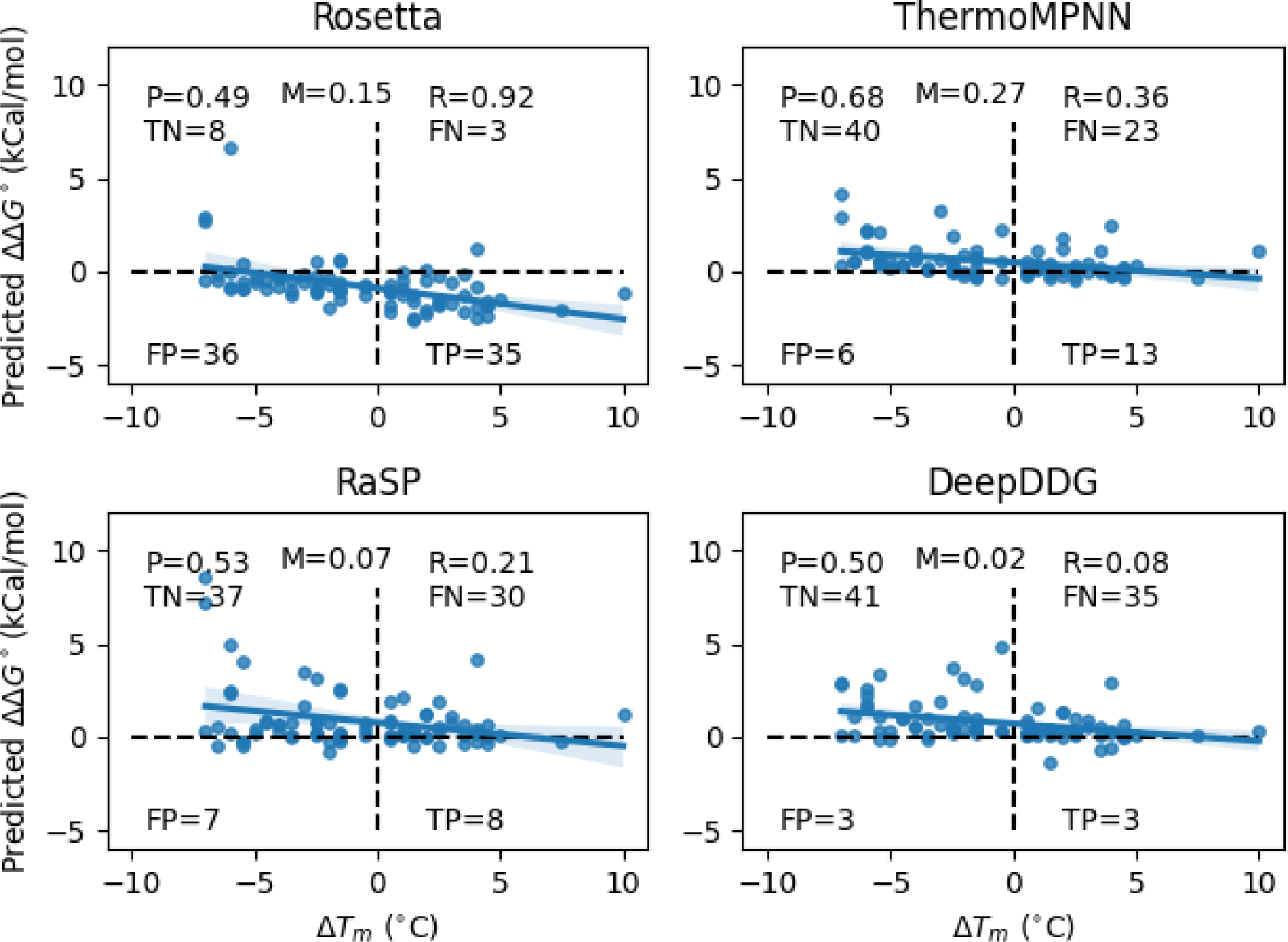
Precision (P), Recall (R), and Mathews correlation coefficient (M) for agreement of computationally predicted change in free energy of unfolding (ΔΔG^0^) with experimental data (ΔT_m_). ThermoMPNN exhibited the highest precision and MCC. However, in terms of recall, Rosetta achieved the highest performance.

### 3.3 Average change in T_m_ of the predicted stabilizing mutations

To further evaluate the accuracy of the computational predictors on their prediction of stabilizing mutations, we analyzed the average ΔT_m_ of the predicted stabilizing mutations (Figure 4A). We randomly sampled 25%, 50%, 75%, and 90% of the experimental data. For each of these chosen sets of mutants, we analyzed the predictions by the ΔΔG^0^ predictors, obtained the subset of mutations predicted to be stabilizing, and calculated their average experimental ΔT_m_ values (Equation 8). This average ΔT_m_ indicates the average change in melting temperature that is expected when these predicted stabilizing mutations are individually incorporated in the wild-type protein.

**Figure 4.**
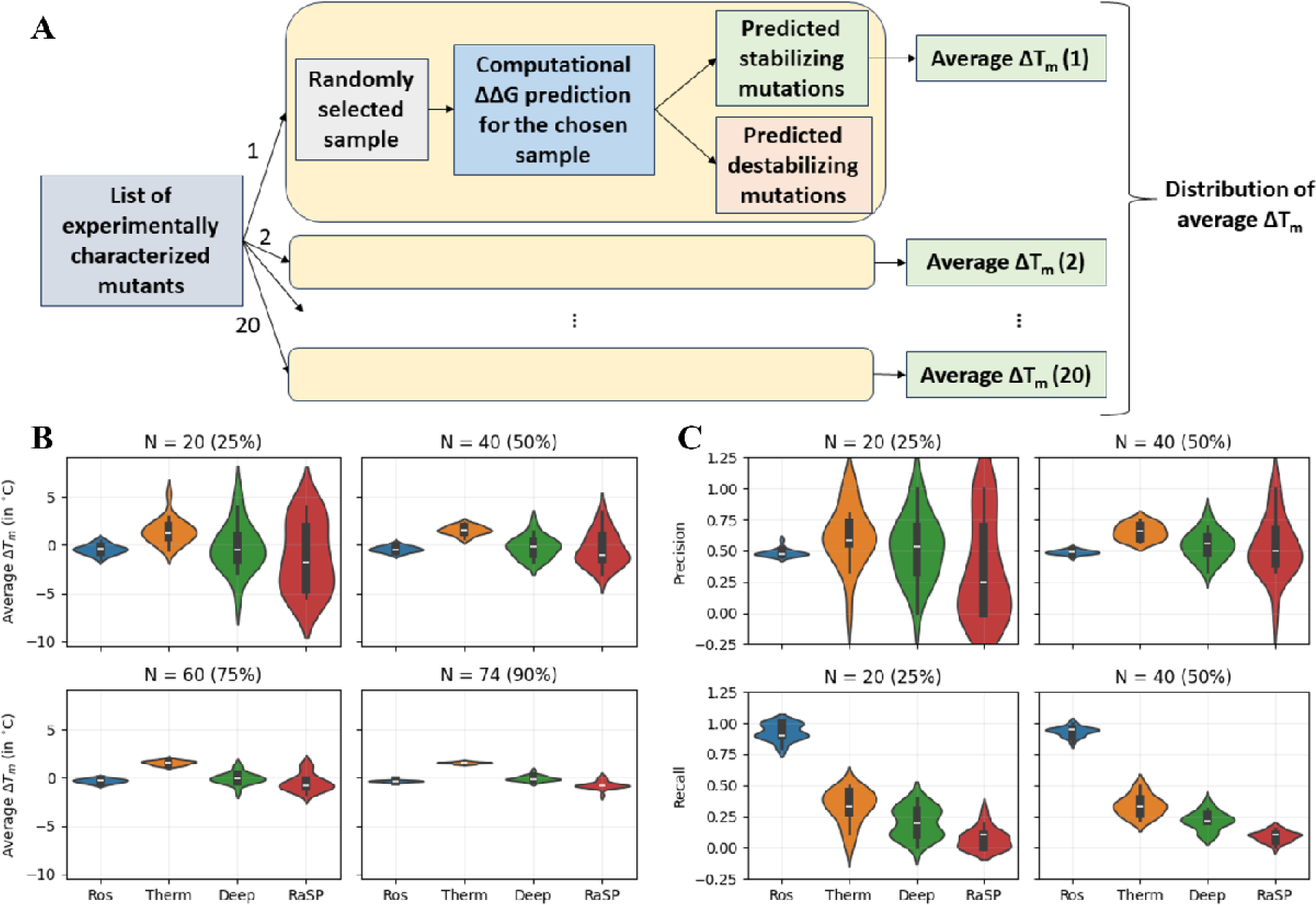
Distribution of the average ΔT_m_ of the predicted stabilizing mutations. (A) Schematic depicting the process of sampling the data to obtain the average ΔT_m_ values. (B) Average ΔT_m_ distribution. From the experimental dataset, 20, 40, 60, and 74 mutants were randomly sampled and the average ΔT_m_ (Equation 8) was calculated for the subset of predicted stabilizing mutants. This was done for 20 iterations and the distribution of the average ΔT_m_ of the mutations predicted to be stabilizing by Rosetta (Ros), ThermoMPNN (Therm), DeepDDG (Deep), and RaSP is shown. The median of the distribution is the highest for mutations predicted to be stabilizing by ThermoMPNN. The mean of the distributions for Rosetta, DeepDDG, and RaSP are all negative, indicating that the majority of the predicted stabilizing mutations were not stabilizing in experiments. When the average is taken for more points (with increasing N), the distributions become narrower but the trends remain the same. (C) Distribution of precision and recall values. We sampled either 25% or 50% of the data 20 times and analyzed the distributions of the precision and recall values. ThermoMPNN predictions had the highest precision, while the Rosetta predictions had the highest recall.

We repeated the above process 20 times and analyzed the distribution of the 20 average ΔT_m_ values (Figure 4B). We found that the average ΔT_m_ values were highest for ThermoMPNN, confirming that it is the best among the chosen computational predictors at identifying stabilizing mutations. We also note that the range of average ΔT_m_ values is narrow for Rosetta and ThermoMPNN while it is wide for DeepDDG and RaSP when 25% of the data is sampled. As the percentage of sampled data increased, all the distributions became narrower, but the trends among the predictors remained consistent.

We sampled 25% and 50% of the data 20 times and analyzed the distribution of the precision and recall values (Figure 4C). The ThermoMPNN predictions had the highest precision, while the Rosetta predictions had the highest recall.

### 3.4 Logistic regression models for identifying stabilizing mutations

We trained logistic regression models to predict the stability change associated with point mutations incorporating values of MFI from yeast surface display experiments as well as computational predictors. We ensured that we had a balanced set of stabilizing and destabilizing mutants in both our training and test sets. First, we trained univariate logistic regression models on MFI values obtained either in the background of the wildtype (MFI_wt_), for different PIMs (MFI_PIM_), or the average of the MFI values for the three PIM libraries (MFI_avg_). Next, we combined the MFI values with each of the computational predictors and developed bivariate regression models (models with two features). We also included residue depth^38–40^ as a feature along with the MFI values, but it did not improve the precision, recall, or average ΔT_m_ of the predicted stabilizing mutations (Supplementary Figure 1).

Univariate and bivariate logistic regression models have two and three parameters, respectively. The training set must have a population that is ideally at least ten times the number of parameters^35^, which is 20 mutants. This corresponds to about 25% of our experimental dataset. Therefore, we trained models with 25% of the data and tested it on the remaining 75% (Figure 5A). To understand the effect of training on a larger set, we also designed another set of models by training on 50% of the data and tested their precision on the remaining 50%.

**Figure 5.**
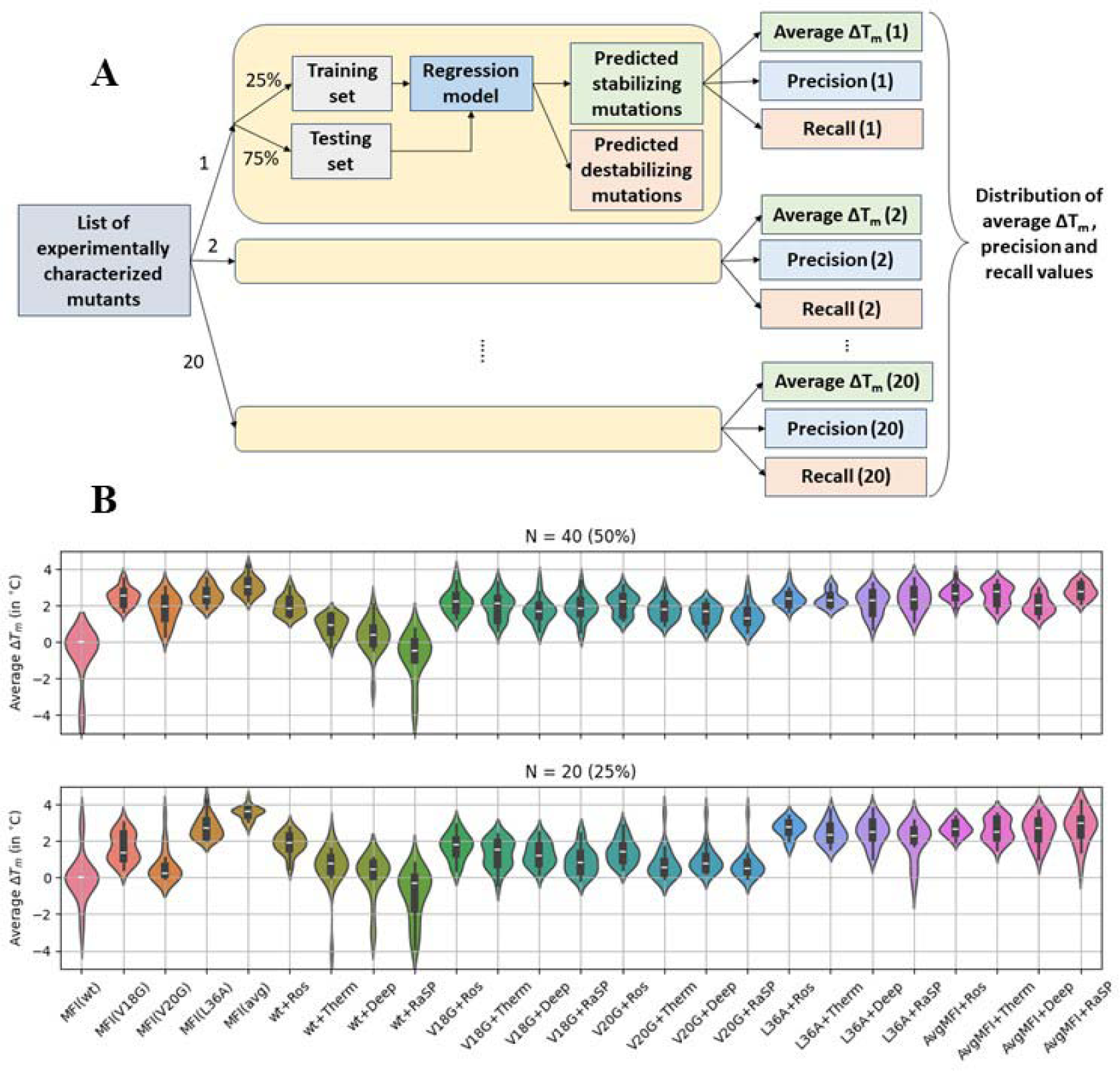
Distribution of the average ΔT_m_ of the predicted stabilizing mutations for different training sets and computational predictors. (A) Schematic representation of the process of splitting the data into train and test sets and calculating the average ΔT_m_ and precision values. (B) The model trained on MFI_avg_ had the highest average ΔT_m_, indicating that most of the predicted stabilizing mutants are stabilizing in experiments. A comparison of predictions from the model trained exclusively on MFI_wt_ with the model trained on both the Rosetta predictions and MFI_wt_ indicates that the latter shows improved predictions. For models trained on MFIs obtained from the individual PIM libraries or the MFI_avg_, including predictions from computational predictors did not result in any improvement in the prediction of stabilizing mutations. The trends are similar irrespective of whether we train the models on 50% (with 40 mutants) or 25% (with 20 mutants) of the experimental data.

Every regression model was trained 20 times to avoid any bias that could arise from the way the data was split into train and test sets. For each set of predicted stabilizing mutations from each iteration, the average experimental ΔT_m_ was calculated (Equation 8). The distribution of this average for the 20 iterations was plotted (Figure 5B). The model trained on MFI_avg_ had the highest average ΔT_m_, indicating that the average MFI from multiple PIM libraries is the best metric to predict stabilizing mutations.

The model trained on MFI_wt_ failed to accurately predict stabilizing mutations. This is attributed to the high stability and well-folded nature of the wild-type protein. Therefore, mutations that enhance the stability of wild-type CcdB are unlikely to further enhance the MFI of binding to its cognate target, GyrA14; only 22% of mutants have an MFI value at least 10% greater than the wild-type. This issue is overcome by incorporating a PIM into the library. Since the PIM has a significantly reduced value of MFI, it is now straightforward to screen for suppressors which enhance the MFI of binding, relative to the corresponding value for the PIM^7^. The best predictions were obtained using the MFI_avg_ values which incorporate data from multiple PIM libraries. Since each PIM library is made by incorporating the PIM mutation into each member of the original site saturation library by a single site-directed mutagenesis reaction, creation and screening of these libraries is straightforward and multiple libraries can be simultaneously screened and sequenced^7^.

Having developed models that used only the high-throughput experimental data as the input, we next built models that used both the high-throughput experimental data and the computational predictions as inputs. This was done to understand if adding the computational predictions to models that already use high-throughput experimental data improves their accuracy. Combining the MFI_wt_ values with the Rosetta predictions led to an improvement in predicting stabilizing mutations as seen from the increase in the average ΔT_m_. For models trained on MFI_PIM_ or MFI_avg_, including predictions from any computational predictor did not result in an improvement in the prediction of stabilizing mutations. The distribution trends remained the same irrespective of whether the models were trained on 50% or 25% of the data.

### 3.5 Precision and recall of the regression models

For an experimentalist, the making, purifying, and characterizing of individual mutants of a protein is laborious. Hence it is desirable that computational predictors have high precision so that minimal numbers of putative stabilizing mutations need to be made experimentally. We therefore analyzed the distributions of the precision values for various predictive models (Figure 6A). We also calculated the recall values to understand what fraction of the actual stabilizing values were being identified by our models (Figure 6B). The model trained on the values of MFI_avg_ had the highest precision in predicting stabilizing mutations while the model trained on MFI_wt_ had the lowest precision. The MFI_avg_ model had a precision of about 0.7. When Rosetta predictions were added to MFI_wt_, the model precision improved to 0.75 and the recall improved to 0.7 suggesting that in cases where the library contains very few mutations with significantly higher binding MFI than the corresponding MFI for WT, input from computational predictors is useful. As discussed above this is because once the WT protein crosses a threshold level of stability and is well folded, further increases in stability do not translate into enhanced protein levels and consequently enhanced binding. For models trained on MFI_PIM_, adding information from computational predictors of ΔΔG^0^ did not result in any improvement in the precision, but there was a small increase in the model recall. The distributions followed the same trend when the models were trained on 50% or 25% of the data.

**Figure 6.**
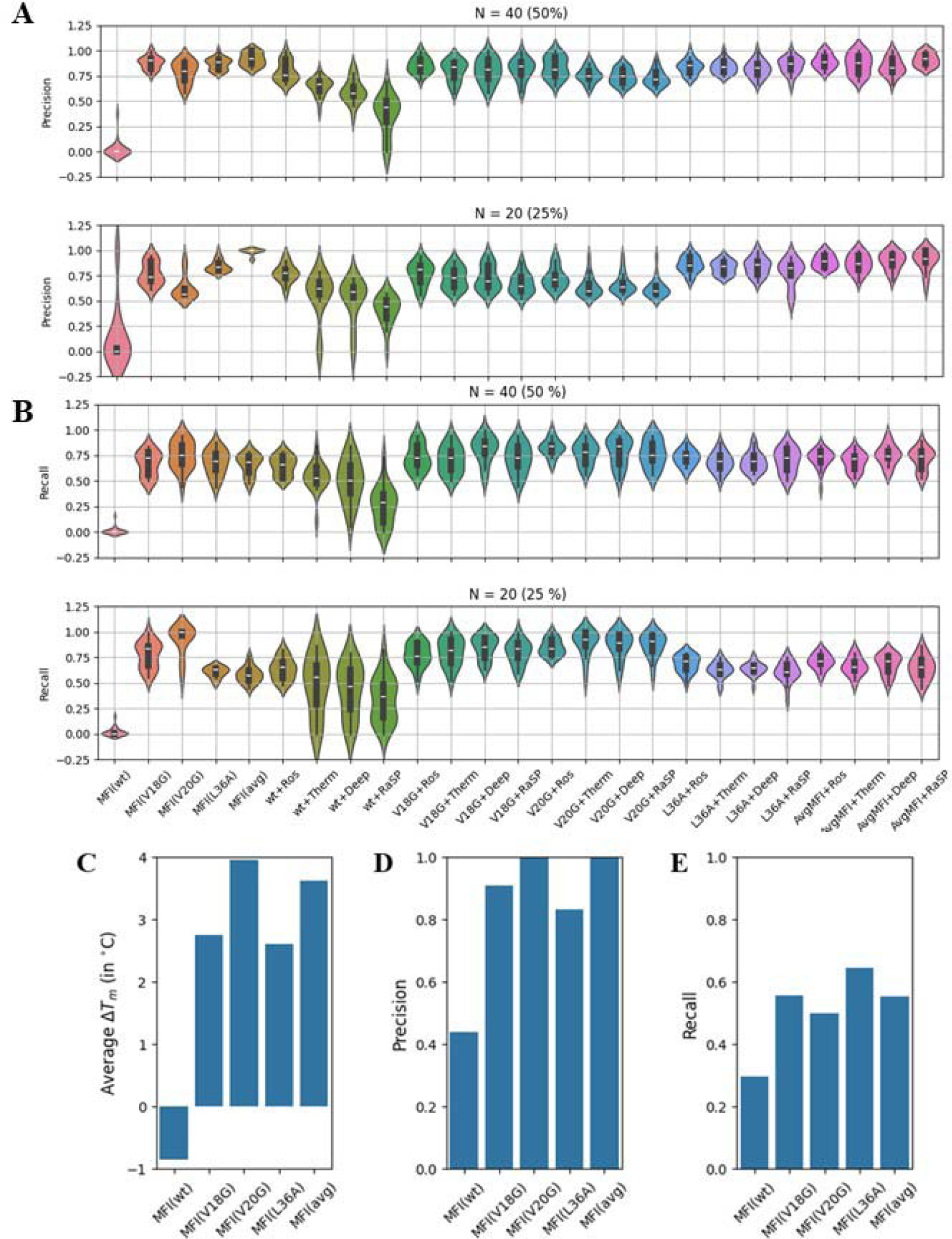
Distribution of precision and recall values for different training datasets and computational predictors. (A-B) The model trained on MFI_avg_ has the highest precision in predicting stabilizing mutations. This model had a recall of 0.7. When ΔΔG^0^ predictions from Rosetta were added to the model trained on MFI_wt_, the resulting model had a significant improvement in both precision and recall. For models trained on MFIs obtained from PIMs, adding data from computational predictors did not result in any increase in the model’s precision but there was a small increase in the recall. The trends are similar irrespective of whether we train the models on 50% or 25% of the data. (C-E) We also predicted stabilizing mutations from the distribution of the MFI values without using the *in vitro* measured mutant thermal stabilities. Mutants with Z-scores > 1 were predicted to be stabilizing. The average ΔT_m_ and precision were highest for mutants predicted using Z-scores from the MFI values from the V20G library and MFI_avg_ and lowest for MFI_wt_.

### 3.6 Identifying stabilizing mutations from mutational binding data alone

Given the high precision and recall for stabilized mutant identification in the model trained on values of MFI_avg_, we next examined if it was possible to infer stabilizing mutations simply by examining the distribution of MFI values in the various libraries, without the requirement for training on mutant thermal stability measured *in vitro*. By choosing mutants with a Z-score (Equation 9) greater than 1, it was possible to identify stabilizing mutations without any training in all libraries, though expectedly, performance was poorest for the WT library. For the suppressor libraries, the largest values of ΔT_m_ were observed using MFI values from the V20G or MFI_avg_ (Figure 6C) and encouragingly, these were similar to corresponding values seen using logistic regression on the MFI_avg_ values.

The methodology used for the high-throughput binding experiment has the following limitations. It requires a conformation-specific ligand to distinguish between the unfolded (destabilized) PIM and folded (stabilized) PIM-suppressor pair. Proteins with N-linked glycosylation sites undergo hyperglycosylation when expressed on the yeast cell surface. In case this interferes with ligand binding it might be necessary to either mutate these sites or express the proteins on the mammalian cell surface. While mammalian cells have a more stringent quality control system than yeast, mammalian cell surface display is technically more difficult to carry out than yeast surface display and for large library sizes it is challenging to obtain the required diversity ^41^.

### 3.7 Assessing ThermoMPNN and LLM predicted mutations for Neuraminidase

ThermoMPNN is trained to predict ΔΔG^0^ for monomeric proteins. However, given its successful predictions for homodimeric CcdB, it would be useful to further evaluate its performance on a more complex target, Neuraminidase (NA), a homotetrameric protein. After analyzing the ThermoMPNN predictions for NA, we observed over 1200 mutations with a predicted ΔΔG^0^ < 0 with 82.5% of them being exposed (with residue depth < 5Å). Considering the impracticality of evaluating such a vast number of mutants *in vitro*, we selected five exposed residues showing the lowest ΔΔG^0^ values for experimental validation.

To enhance the reliability of the predictions, we also experimentally tested six additional mutants suggested by both ThermoMPNN and a protein evolution algorithm that uses the ESM large language model (LLM) trained solely on protein sequences^42^. Although this algorithm was trained to predict evolutionarily plausible mutations, most of the predicted mutations were also found to improve the thermostability of the designed antibodies^43^. Therefore, we experimentally assessed the stability of a total of 11 mutations relative to wild-type NA. For this purpose, full-length wild-type and NA mutants were expressed on the HEK cell surface, and evaluated for stability by measuring their binding to 1G01, 2H08, and FNI17 antibodies (Figure 7). However, none of the tested mutants showed better binding than the wild-type. These results suggest that the current version of ThermoMPNN needs to be further optimized for ΔΔG^0^ predictions of complex oligomeric proteins.

**Figure 7.**
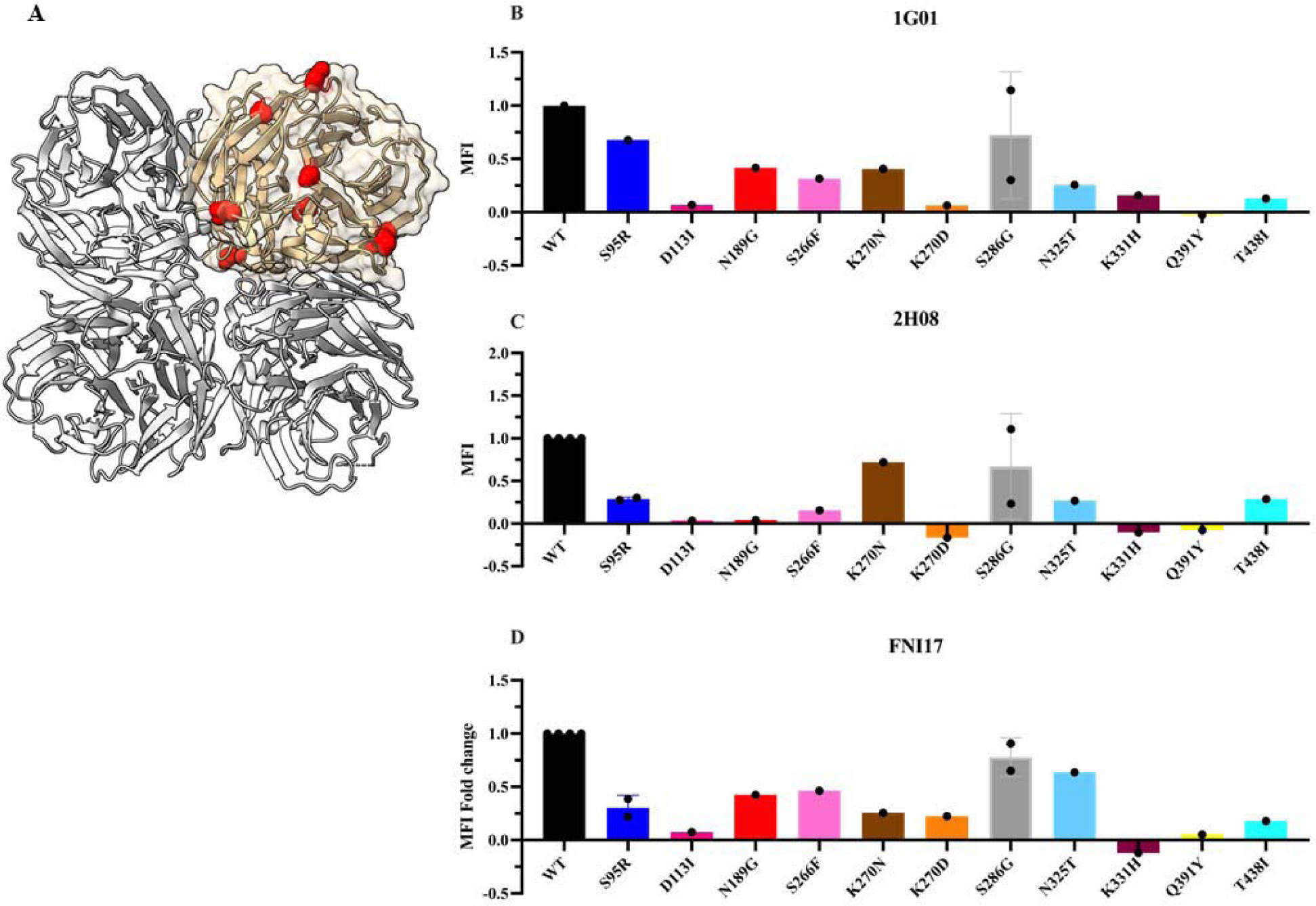
Binding profiles of NA mutants relative to the wild type (WT). (A) Predicted stabilizing NA mutants mapped on the structure of NA (PDB: 7U2Q), visualized using ChimeraX^44^. (B-D) Binding profiles. HEK-293 cells expressing the wild-type and mutant NA constructs were incubated with NA-specific antibodies (B)1G01, (C) 2H08, and (D) FNI17, and their binding profiles were analyzed using FACS. The fold change in MFI was calculated as the ratio of the MFI of the mutant to the MFI of the wild type. None of the ThermoMPNN predicted stabilizing mutants that we tested showed better binding than the wild-type NA, suggesting that they are unlikely to be stabilizing.

## Conclusions

Predicting stabilizing mutations in proteins using computational methods is challenging. For the CcdB protein, we found that the best-performing computational predictor, ThermoMPNN, had a precision of 0.68. The average increase in thermal melting temperature of the predicted stabilizing mutants was about 1°C. To further improve predictions of stabilizing mutations, we trained logistic regression models on experimental values of MFI of binding. The prediction of stabilizing mutations from MFI values in the wild-type library was inaccurate, however, this could be improved by combining these results with those from computational predictors. In contrast, the model trained on the average MFI from three PIM libraries had the highest precision. The average increase in T_m_ of the predicted stabilizing mutants by this model was about 3-4°C. We also demonstrate that it is possible to accurately infer stabilizing mutations simply by examining the distribution of MFI values in the various libraries, without the requirement for training on mutant thermal stability measured *in vitro*. Additionally, we used ThermoMPNN to predict ΔΔG^0^ for all single mutants of the Influenza surface glycoprotein, NA, and measured the binding of predicted stabilizing mutants relative to the wild-type. However, none of the tested mutants showed better binding than the wild-type, indicating the need to develop better predictors for stabilizing mutations in complex oligomeric proteins. Our analysis of the CcdB protein shows that MFI binding values obtained from carefully designed experiments in conjunction with or without limited stability data, can be used to develop protein-specific predictions of stabilizing mutations. Since MFI or alternative measures of binding data can be obtained from high-throughput experiments, predictions of stabilizing mutations can be made rapidly and accurately. Once such data is available for multiple systems, it should be also possible to find an optimal way of combining such experimental binding data with computational stability predictions, to further enhance stability predictions for a given protein of interest. In the longer term, the increasing availability of such data can in turn be used to train computational models to predict stabilizing mutations without recourse to prior experiments for the specific protein of interest.

## Data availability statement

All data are available in the main text

## Acknowledgments

This work was funded in part by a grant to RV from the Science and Engineering Research Board, Government of India (CRG/2022/004425) and through the ENDFLU project. The ENDFLU project has received funding from the European Union’s Horizon 2020 research and innovation program under grant agreement No. 874650 and from the Department of Biotechnology, Ministry of Science and Technology, Government of India (BT/IN/EU-INF/15/RV/19-20). Funding for infrastructural support was from DST FIST, UGC Center for Advanced Study, MHRD, and the DBT IISc partnership program. NM acknowledges the Prime Minister Research Fellowship for her fellowship (PM/MHRD-20-17303.03). RV is a JC Bose Fellow of DST. We thank Prof. Nagasuma Chandra (Department of Biochemistry, Indian Institute of Science) for generously letting us use her Intel Xeon cluster for this work. Dr. Shahbaz Ahmed is acknowledged for the data from deep mutational scans and thermal stability experiments.

## Author contributions

SG carried out the analysis and drafted the manuscript. NM did experimental validation of NA mutants. DN, AB, and RA carried out Rosetta calculations for CcdB. SG and AG trained the regression models. RV designed the study and supervised the analysis. SG and RV wrote the manuscript with contributions from all co-authors. All authors have read and agreed to the submitted version of the manuscript.

